# Modelling the gastrointestinal carriage of *Klebsiella pneumoniae* infections

**DOI:** 10.1101/2022.10.03.510744

**Authors:** Ricardo Calderon-Gonzalez, Alix Lee, Guillermo Lopez-Campos, Steven J. Hancock, Joana Sa-Pessoa, Amy Dumigan, Ronan McMullan, Eric L. Campbell, Jose A. Bengoechea

## Abstract

*Klebsiella pneumoniae* is a leading cause of nosocomial and community acquired infections, making *K. pneumoniae* the second pathogen associated with the most deaths attributed to any antibiotic resistant infection. *K. pneumoniae* colonises the nasopharynx and the gastrointestinal tract in an asymptomatic manner without dissemination to other tissues; importantly gastrointestinal colonisation is a requisite for infection. Our understanding of *K. pneumoniae* colonisation is still based on interrogating mouse models in which animals are pre-treated with antibiotics to disturb the colonisation resistance imposed by the gut microbiome. In these models, infection disseminates to other tissues. Here, we report a murine model to allow for the study of the gastrointestinal colonisation of *K. pneumoniae* without tissue dissemination. Hypervirulent and antibiotic resistant strains stably colonise the gastrointestinal tract of in an inbred mouse population without antibiotic treatment. The small intestine is the primary site of colonisation followed by a transition to the colon over time without dissemination to other tissues. Our model also mimics the disease dynamics of metastatic *K. pneumoniae* strains able to disseminate from the gastrointestinal tract to other sterile sites. Colonisation is associated with mild to moderate histopathology, no significant inflammation, and no effect on the richness of the microbiome. Our model recapitulates the clinical scenario in which antibiotic treatment disturbs the colonisation of *K. pneumoniae* resulting in dissemination to other tissues. Finally, we establish that the capsule polysaccharide is necessary for the colonisation of the large intestine whereas the type VI secretion system contributes to colonisation across the gastrointestinal tract.

**IMPORTANCE:** *Klebsiella pneumoniae* is one of the pathogens sweeping the World in the antibiotic resistance pandemic. *Klebsiella* colonises the nasopharynx and the gut of healthy subjects in an asymptomatic manner, being gut colonisation a requisite for infection. This makes essential to understand the gastrointestinal carriage to prevent *Klebsiella* infections. Current research models rely on the perturbation of the gut microbiome by antibiotics, resulting in an invasive infection. Here, we report a new model of *K. pneumoniae* gut colonisation that recapitulates key features of the asymptomatic human gastrointestinal tract colonisation. In our model, there is no need to disturb the microbiota to achieve stable colonization without dissemination to other tissues. Our model recapitulates the clinical scenario in which antibiotic treatment triggers invasive infection. We envision our model will be an excellent platform to test therapeutics to eliminate *Klebsiella* asymptomatic colonisation, and to investigate factors enhancing colonisation and invasive infections.

## INTRODUCTION

*Klebsiella pneumoniae* is one of the pathogens sweeping the World in the antibiotic resistance pandemic. Recent analysis of the global burden of antibiotic resistant infections revealed that more than 250,000 deaths are associated with *K. pneumoniae* infections (1), making *K. pneumoniae* the second pathogen associated with the most deaths attributed to any antibiotic resistant infection. Particularly, carbapenem-resistant *K pneumoniae*, and third-generation cephalosporin-resistant *K. pneumoniae* account for at least 100,000 deaths (1). In addition, *Klebsiella* species are a known reservoir for antibiotic resistant genes, which can spread to other Gram-negative bacteria. Not surprisingly, the World Health Organization has singled out *K. pneumoniae* as an urgent threat to human health.

*K. pneumoniae* is a member of the human microbiota found in the mouth, nares, and skin, and the gastrointestinal tract. In Western countries up to 5% of healthy humans from the community are nasopharyngeally colonised with *K. pneumoniae*, and this percentage may increase to 30% in Asian countries (2). In healthy individuals in Western countries, the colonisation of the gastrointestinal tract ranges from 5% to 35%, and this can reach up to 60-70% in Asian countries (2). Importantly, clinical studies demonstrate that gastrointestinal colonisation is a requisite for infection (3, 4). This colonisation is asymptomatic and, therefore, these healthy subjects could act as silent carriers from which *K. pneumoniae* may cause disease when the host is compromised or act as a source of transmission. A point of concern is the colonisation by so-called hypervirulent *K. pneumoniae* (2). These strains are prevalent in Asia, and they have the capacity for metastatic spread. Alarmingly, there are reports of hypervirulent strains becoming multidrug resistant (5-8).

Significant knowledge gaps exit on the host factors influencing gastrointestinal colonisation, although recent data indicates that age, and alcohol consumption are associated with increased colonisation (9). In addition, antibiotic treatment seems to predispose individuals to colonisation (9) and, in the clinical setting, may result in dissemination of *K. pneumoniae* from the gastrointestinal tract to other tissues resulting in sepsis and other life-threating complications (3, 4, 10). These observations suggest that the commensal gut microbiota provides a barrier to *K. pneumoniae* colonisation. Indeed, a number of studies in mice demonstrate that antibiotic pre-treatment facilitates *K. pneumoniae* colonisation (11).

The vast majority of our knowledge on *K. pneumoniae*-associated gastrointestinal colonisation comes from mouse models in which the animals are pre-treated with antibiotics (11), resulting in an infection that disseminates to other tissues. These models do not recapitulate the asymptomatic colonisation of human healthy subjects harbouring an undisturbed microbiome. Furthermore, these models also present limitations to identify *K. pneumoniae* factors implicated in the gut colonisation because the colonisation resistance imposed by the microbiome is impaired as well as there may be changes in the immune response following infection. Undoubtedly, these changes in the gut microenvironment affect the host-*K. pneumoniae* interface, resulting in changes in the factors deployed by the pathogen to ensure colonisation.

Here, we describe a murine model to allow for the study of the gastrointestinal colonisation of *K. pneumoniae*. We demonstrate that *K. pneumoniae* can stably colonise the gastrointestinal tract of in an inbred mouse population without antibiotic pre-treatment. We characterize the colonisation dynamics by *K. pneumoniae*, and show that antibiotic treatment triggers the dissemination of the infection. Finally, we establish the role of the capsule polysaccharide (CPS) and implicate the type VI secretion system (T6SS) in the colonisation of the gastrointestinal tract.

## RESULTS

### Establishing *K. pneumoniae* colonisation of the murine intestinal tract

We sought to establish a murine model of gastrointestinal tract colonisation by *K. pneumoniae* that would mimic the asymptomatic gastrointestinal colonization of healthy individuals without dissemination to other tissues. Previous studies have used oral gavage and pre-treatment with antibiotics to disrupt the colonisation resistance imposed by the microbiome (11). We first tested the ability of *K. pneumoniae* CIP52.145 (hereafter Kp52145) to colonize the gastrointestinal tract of adult mice (8-9 weeks) of both sexes without antibiotic treatment. This strain belongs to the *K. pneumoniae* KpI group and it encodes all virulence functions associated with invasive community-acquired disease in humans (12, 13). A pilot experiment using oral gavage and a dose of 10^8^ CFU per mouse did not result in colonisation as measure by the presence of bacteria in the feces by plating on SCAI medium (14) 10 days post infection. Fecal shedding is a widely used substitute for colonization density in the gastrointestinal tract. SCAI medium consists of Simmons Citrate agar and inositol. *Klebsiella spp* grow as yellow colonies, whereas, apart from few *Enterobacter* and *Citrobacter* strains, other typical *Enterobactericeae* found in the gut grow as blue colonies due to citrate utilization (14). We reasoned that the lack of detection of colonisation could be the result of low numbers of bacteria reaching the gastrointestinal tract. One of the innate barriers protecting the gastrointestinal tract from infections is the gastric acid of the stomach. Therefore, we hypothesized that colonisation would be facilitated if the gastric acid of the stomach is quenched. Indeed, treatment of mice with sodium bicarbonate 5 min before oral gavage with *K. pneumoniae* resulted in gastrointestinal colonisation by Kp52145 (Fig S1A).

Having established that sodium bicarbonate treatment facilitated colonisation, we next sought to determine the minimal dose of *K. pneumoniae* resulting in a reproducible gastrointestinal colonisation. Mice were infected with four bacterial concentrations, and the weight loss, as an indication of health status, was measured daily. Three of the six mice infected with 3×10^8^ CFUs reached the humane threshold to euthanize them within five days post infection. The remaining three mice lose 6% of weight by day eight post inoculation although subsequently they recovered weight reaching the weight before inoculation. Mice infected with 5×10^7^, 1×10^7^ and 5×10^6^ also lost weight which peaked at day six post infection before starting to gain weight. After, we observed no more than 5% difference between infected and non-infected mice (Fig 1A). At the end of the experiment, day twelve post infection, mice were euthanized and bacterial loads in the intestine were quantified by plating. Kp52145 was detected in the small and large intestine (Fig 1B and 1C). No significant differences between doses were found in the small intestine (Fig 1B). In contrast, mice infected with 1×10^7^ had higher bacterial loads in the large intestine than those infected with either 5×10^7^ or 5×10^6^ (Fig 1C). We next determined the distribution of bacteria along the gastrointestinal tract as percentage. Our results showed that approximately 60% of bacteria were found in the small intestine and without significant differences between doses (Fig 1D). Interestingly, the bacterial loads in faeces were at least ten time lower than those found by plating the tissue (Fig S1B), suggesting that faecal shedding is not a reliable surrogate of gastrointestinal gut colonisation in our model. Taken together, our data suggests that quenching the gastric acid facilitates the colonisation of the gastrointestinal tract by *K. pneumoniae* without need for antibiotic pre-treatment. We chose 1×10^7^ as the dose for subsequent experiments considering its minimal effect on the health of the animals and the levels of bacterial loads across the tissue.

**Figure 1.**
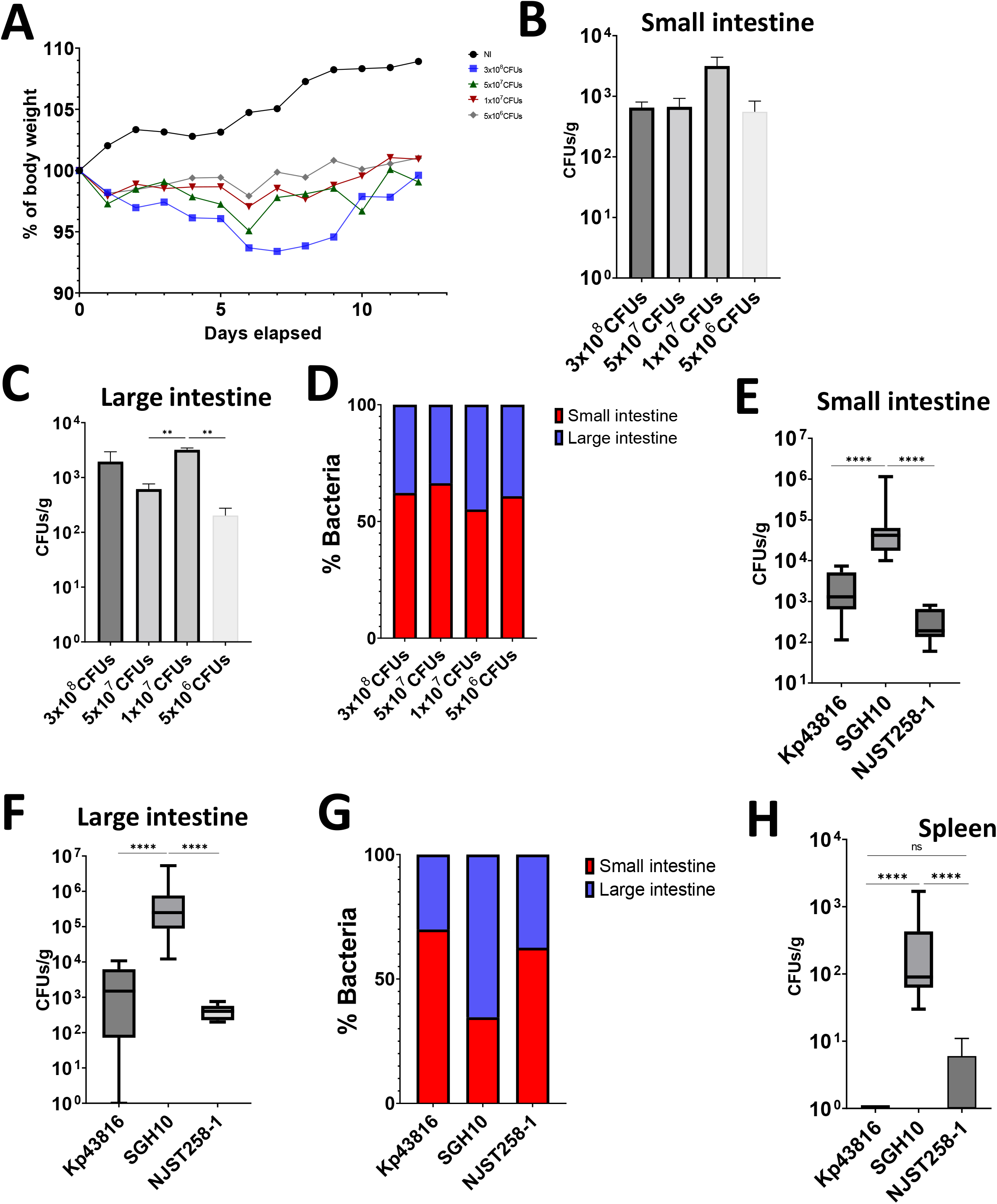
*K. pneumoniae* colonises the gut of immunocompetent mice without need for antibiotic pre-treatment. **A**. Weight loss of mice infected with different doses of Kp52145 over 12 days. 4-6 mice were included in each group except in the group infected with 3×10^8^ CFUs because three mice died within five days post infection. **B**. and **C**. CFUs per gr of small (B) and large intestine (C) of the mice infected in panel A twelve days post infection. **D**. The relative distribution of *K. pneumoniae* across the intestine sections was determined taken as 100% the combined bacterial loads of the small and large intestine sections. **E**. and **F**. Bacterial loads in the small (E) and large (F) intestine of mice infected with Kp43816, SGH10, and NJST258-1. In each group 9-10 mice were infected. **G**. Relative distribution of *K. pneumoniae* strains across the intestine sections. **H**. Bacterial loads in the spleen of infected mice with different *K. pneumoniae* strains twelve days post infection. In all panels, values are presented as the mean ± SD. ****P ≤ 0.0001; ** P ≤ 0.01; ns P > 0.05 for the indicated comparisons determined using one way-ANOVA with Bonferroni contrast for multiple comparisons test.

A feature of *K. pneumoniae* population structure is the genetic diversity between strains (12), therefore, we tested the ability of a set of genetically diverse isolates to colonise the gastrointestinal tract. We chose strain ATCC43816 (hereafter Kp43816), a widely used isolate used to investigate *K. pneumoniae* virulence (15), NJST258-1, an isolate of the carbapenem resistant ST258 epidemic clonal group (16), and SGH10, reference strain of the hypervirulent CG23 clonal group associated with liver abscess infections (17). The three strains colonised the small and large intestine twelve days post inoculation, although the bacterial loads of strain SGH10 were higher than those of the two other strains (Fig 1E and F). Likewise Kp52145, the distribution of Kp43816 and NJST258-1 within the gastrointestinal tract was significantly higher in the small intestine than in the large intestine (Fig 1G). No significant differences between Kp43816 and NJST258-1 were observed. In contrast, the distribution of SGH10 was higher in the large intestine than in the small intestine (Fig 1G). Only in mice infected with SGH10 we observed a significant dissemination to spleen (Fig 1H). Collectively, our results suggest that the gastrointestinal colonisation is a general feature of *K. pneumoniae*. Furthermore, our colonisation model mimics the disease dynamics of metastatic strains such as SGH10 able to disseminate from the gastrointestinal tract to other sterile sites.

### Gastrointestinal dynamics of *K. pneumoniae* colonisation

The differences in bacterial numbers between the small and large intestine led us to explore the colonization dynamics over time in the different sections of the gastrointestinal tract. At three days post infection, bacteria were only detected in the small intestine but not in the cecum or colon (Fig 2A). The bacterial loads in the small intestine remained consistent until the end of the experiment at day twelve (Fig 2A). Colonisation of the cecum was detected first at six days post infection, and it remained constant until the end of the experiment (Fig 2A). However, the bacterial loads in the cecum did not reach those found in the small intestine (Fig 2A). The colonisation of the colon was apparent at day six post infection and reached the levels found in the small intestine at day twelve (Fig 2A). Distribution analyses of the percentage of bacteria in the three sections of the gastrointestinal tract over time further showed the initial colonization of the small intestine with a progressive increase in the number of bacteria colonizing the colon (Fig 2B). The dissemination to other tissues, spleen and lung, was not observed at any time point. Together, these results suggest that the primary site of colonization of the gastrointestinal tract by *K. pneumoniae* is the small intestine followed by a transition to the colon over time but without dissemination to other tissues.

**Figure 2.**
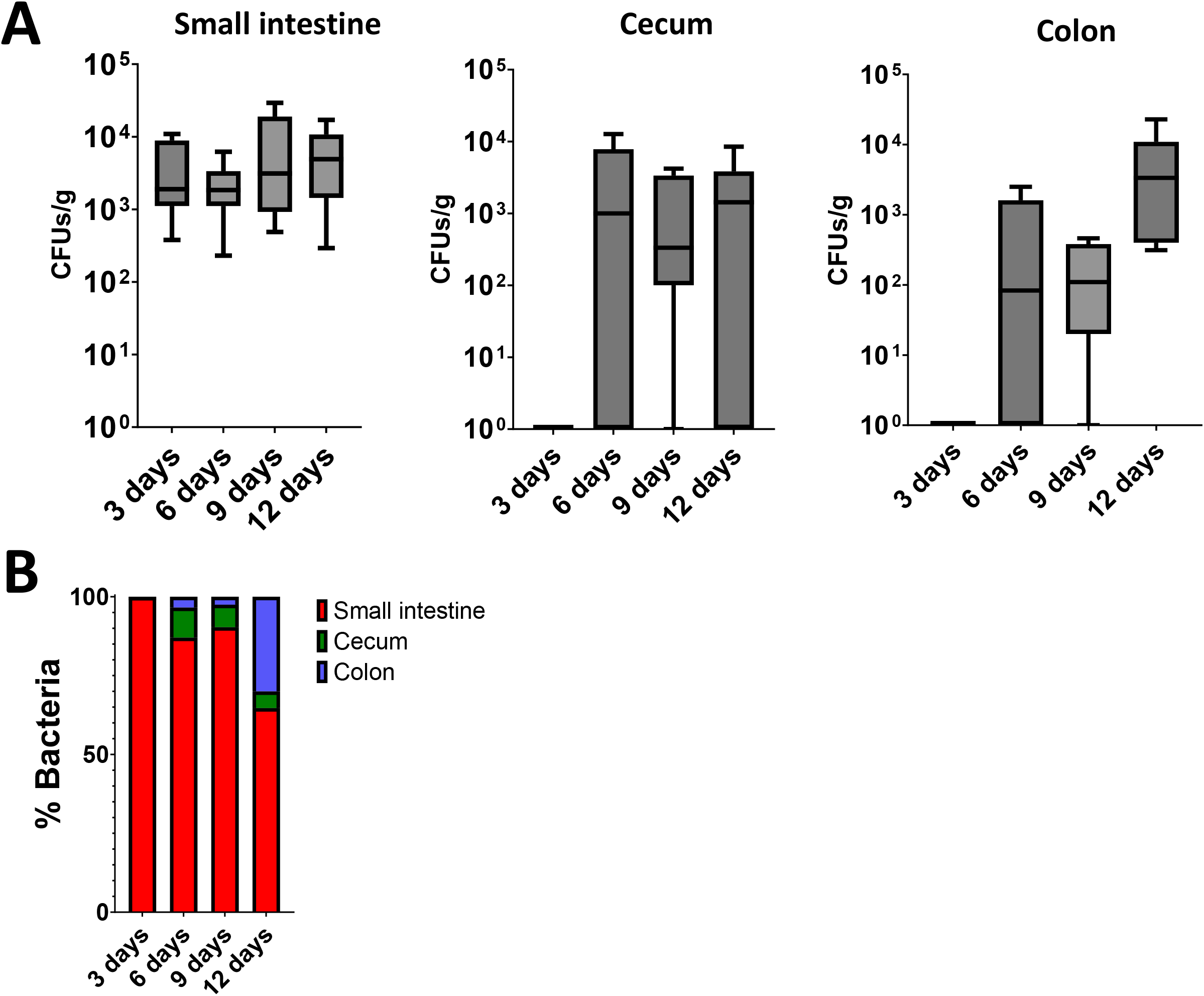
Dynamics of *K. pneumoniae* gut colonisation. **A**. Bacterial loads in the different intestine sections of mice infected with Kp52145 at different days post infections. Values are presented as the mean ± SD. **B**. Relative distribution of Kp52145 across the intestine sections.

We next investigated whether *K. pneumoniae* colonisation is associated with histopathological changes in the gastrointestinal tract. We screened sections for the infiltration of inflammatory cells, presence of submucosal oedema, epithelial damage, or the presence of exudate in the small intestine and colon sections stained with haematoxylin-eosin according to the criteria shown in Table S1 (18). A combined score between 4 and 6 is considered moderate inflammation whereas a score higher than 7 indicates severe inflammation (18). Figure 3 shows mild to moderate epithelial damage, accumulation of exudate, and infiltration of inflammatory cells at day six and twelve post infection in the small intestine of infected mice; although these changes were not significantly different than those of non-infected mice. Similar mild to moderate changes were observed in the colon of infected mice at six and twelve days post infection (Fig 3A and 3B).

**Figure 3.**
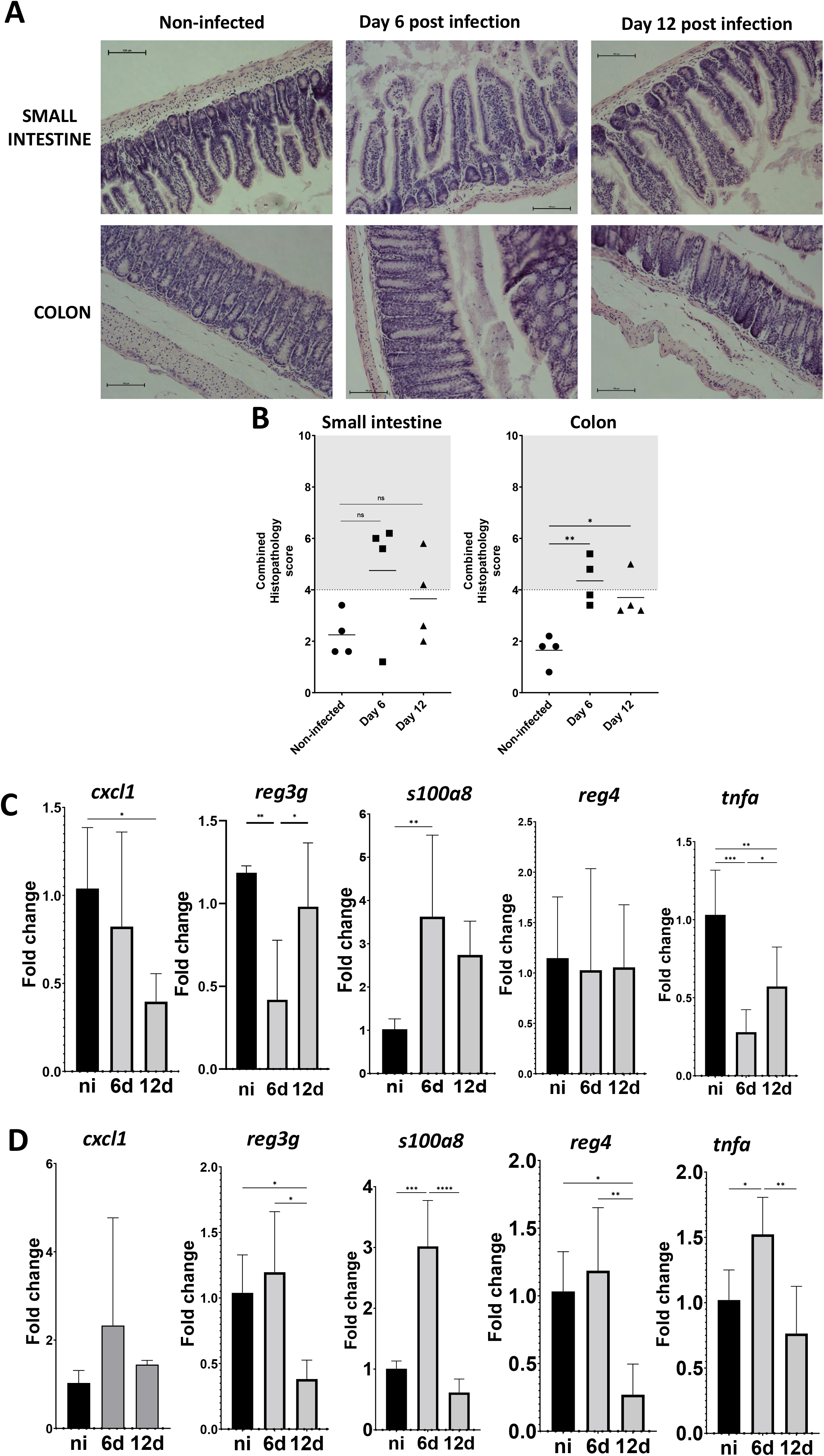
*K. pneumoniae* colonisation does not produce sever tissue damage. **A**. Haematoxylin-eosin staining of tissue at different days post infection with Kp52145. **B**. Quantification of the histopathology changes upon infection. Each dot represents a different mice. Five slides per section of tissue, time point and per mouse were scored. **C**. and **D**. *cxcl1, reg3g, s100o8, reg4* and *tnfa* mRNA levels were assessed by qPCR in the small intestine (C) and the colon (D) of non-infected mice (black bars), and infected mice (grey bars) at six (6d) and twelve (12d) days. In all groups 5-7 mice were analysed. In panel A, the images are representative of four infected mice. Values are presented as the mean ± SD. ****P ≤ 0.0001; ***P≤ 0.001;** P ≤ 0.01; *P ≤ 0.05 for the indicated comparisons determined using one way-ANOVA with Bonferroni contrast for multiple comparisons test. Any other comparison are not significant (P>0.05).

To provide further evidence that the colonisation is not associated with major alterations in the tissue, we next assessed the expression of inflammatory genes associated with host defence against gut pathogens, and inflammation. We detected the upregulation of *s100a8* only at six days post infection in the small intestine and the colon (Fig 3C and 3D). S100a8 is present in intestinal tissue during inflammation, and it is considered as a valuable biomarker of colonic inflammatory conditions (19). Kp52145 colonization did not upregulate the inflammatory marker *cxcl1* in either the small intestine or the colon (Fig 3C and 3D) whereas infection upregulated *tnfa* only six days post infection in the colon (Fig 3C and 3D). Infection did not induce the expression of the regenerating islet-derived family members 4 and 3g (Fig 3C and 3D). The upregulation of Reg proteins is observed in intestinal inflammation, and some of the Reg family members they show antimicrobial activity (20, 21). These results indicate that *K. pneumoniae* gut colonisation does not induce inflammation.

Finally, we sought to determine whether *K. pneumoniae* colonization perturbs the gastrointestinal tract microbiota. Samples were obtained at six and twelve days post infection from the small intestine and the colon, and the intestinal microbiota assessed by 16S rRNA sequencing. No significant differences were observed in the alpha diversity independently on the test used to measure it (Fig S2). At the phylum level, colonisation with Kp52145 did not lead to dramatic changes in the relative abundance (Fig 4A). In these samples we did not detect *K. pneumoniae* 16S rRNA gene sequences, suggesting that *K. pneumoniae* comprises only a minor component of the host intestinal microbiota.

**Figure 4.**
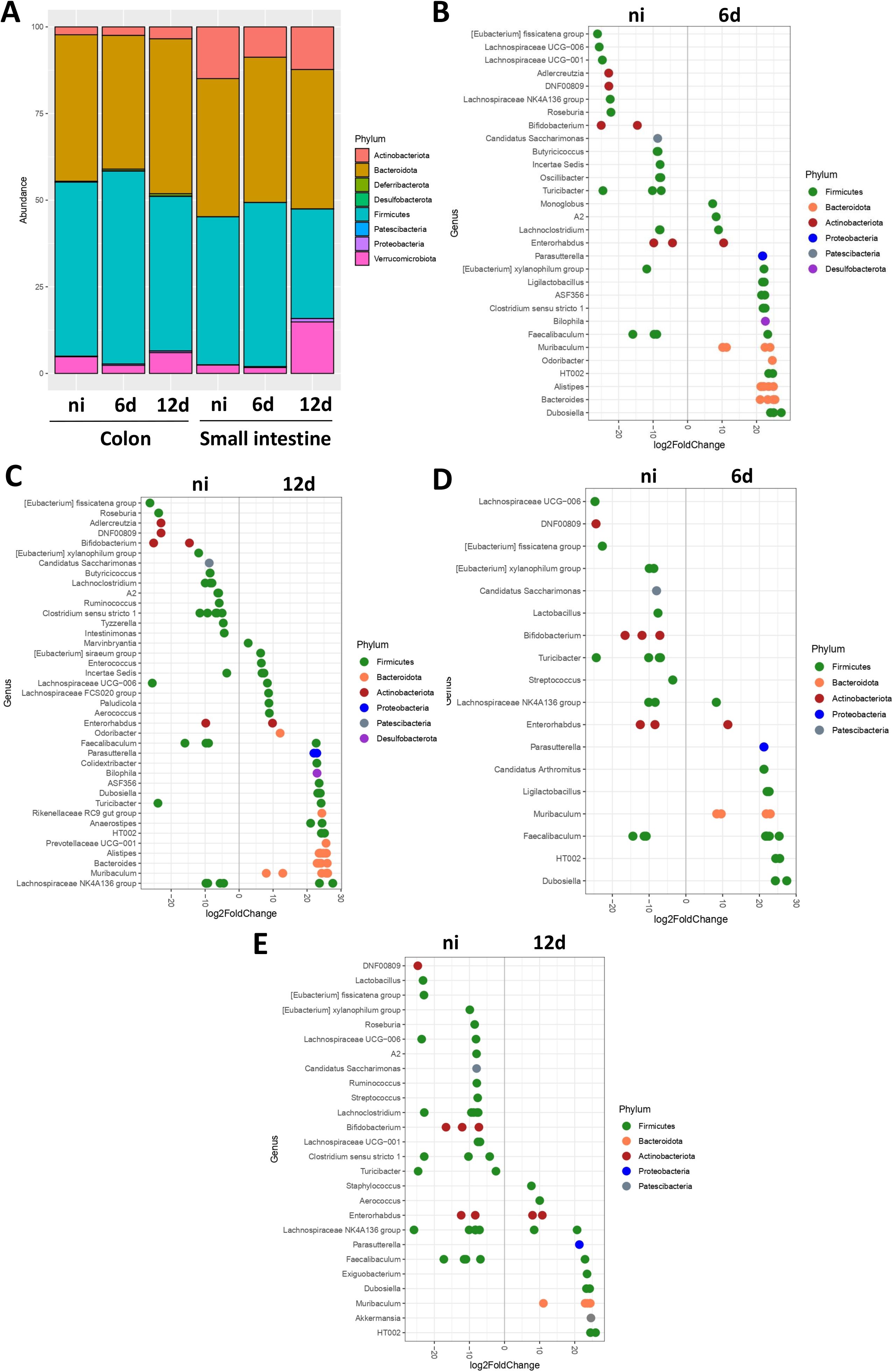
*K. pneumoniae* colonisation has minor effect on the bacterial gut microbiome. **A**. Phyla present in the colon and small intestine of infected mice with Kp52145 were determined by 16S rRNA sequencing. **B** and **C**. Log2 fold change of the genera present in the colon of infected mice at six (B) and twelve (C) days post infection versus the non-infected (ni) mice. **D**. and **E**. Log2 fold change of the genera present in the small intestine of infected mice at six (D) and twelve (E) days post infection versus the non-infected (ni) mice. In all panels, 5-7 mice were analysed in each group.

The colon samples were characterized by the abundance of *Firmicutes* and *Bacteriodes*, and colonisation was associated with a relative increase in the abundance of *Desufolbacteria* and *Proteobacteria* at twelve days post infection. At the genus level, the colonization of the colon at six days post infection resulted in changes within the species of *Actinobacteria*, particularly a decrease in the relative abundance of *Bifidobacterium* and *Adlercreutzia* (Fig 4B). We also detected changes within the species of *Firmicutes*, notably an increase in the relative abundance of *Clostridium* (Fig 4B). The *Proteobacteria Parasutterela*, the *Desulfobacterium Biolophila*, and the *Bacteroides Muribaculum, Prevotella, Alistipes* and *Bacteroides* were characteristic of Kp52145-colonized colon (Fig 4B). Similar findings were obtained when analysing the samples from twelve days post infection except the relative increase in *Clostridium* (Fig 4C). The main phyla comprising the small intestine samples were *Actinobacteria, Bacteroides*, and *Firmicutes* (Fig 4A). Only at twelve days post infection, we detected changes in the microbiota of the small intestine with a relative increase in *Verrrucomicrobiota* and *Proteobacteria*, and a relative decrease in *Firmicutes* (Fig 4A). At the genus level, at six days post infection, conspicuous changes were the decrease in the relative abundance of *Bifidobacterium*, and an increase in the abundance of *Parasutterela* and *Muribaculum* (Fig 4D). Similar changes were observed at twelve days post infection (Fig 4E). Additionally, we observed an increase in the relative abundance of *Akkermansia*, and a shift in the genera of *Firmicutes* in the Kp52145-colonized samples. (Fig 4E). Altogether, these findings demonstrate that although *K. pneumoniae* colonization did not affect the richness of the microbiome, changes were observed at the phylum and genus levels. *K. pneumoniae* colonisation results in shifts in the species of *Firmicutes*, a decrease in the relative abundance of the *Actinobacteria Bifidobacterium*, and an increase in the abundance of the *Proteobacteria Parasutterela*, and of species of *Bacteroides*.

### Antibiotic treatment triggers the dissemination of *K. pneumoniae* from the gastrointestinal tract

In a clinical setting, patients received antibiotic treatment and evidence indicates that this may result in *K. pneumoniae* dissemination from the gut to other tissues (10). Therefore, we sought to establish whether antibiotic treatment has any effect on *K. pneumoniae* that colonises the gastrointestinal tract. Mice were colonised with the ampicillin resistant strain Kp43816, and after six days they received two doses of ampicillin (Fig 5A). Antibiotic treatment resulted in two logs increase in the number of Kp43816 in the small intestine (Fig 5B). Antibiotic treatment also increased dramatically the levels of Kp43816 in the cecum and colon (Fig 5B). Distribution analysis further confirmed that antibiotic treatment shifted the colonization of *K. pneumoniae* from the small to the large intestine (Fig 5C). Antibiotic treatment also caused the dissemination of Kp43816 from the gut to the spleen, liver and lungs (Fig 5D). Altogether, these findings demonstrate that our gut colonisation model recapitulates the clinical scenario in which antibiotic treatment disturbs the colonisation of *K. pneumoniae* resulting in dissemination to other tissues.

**Figure 5.**
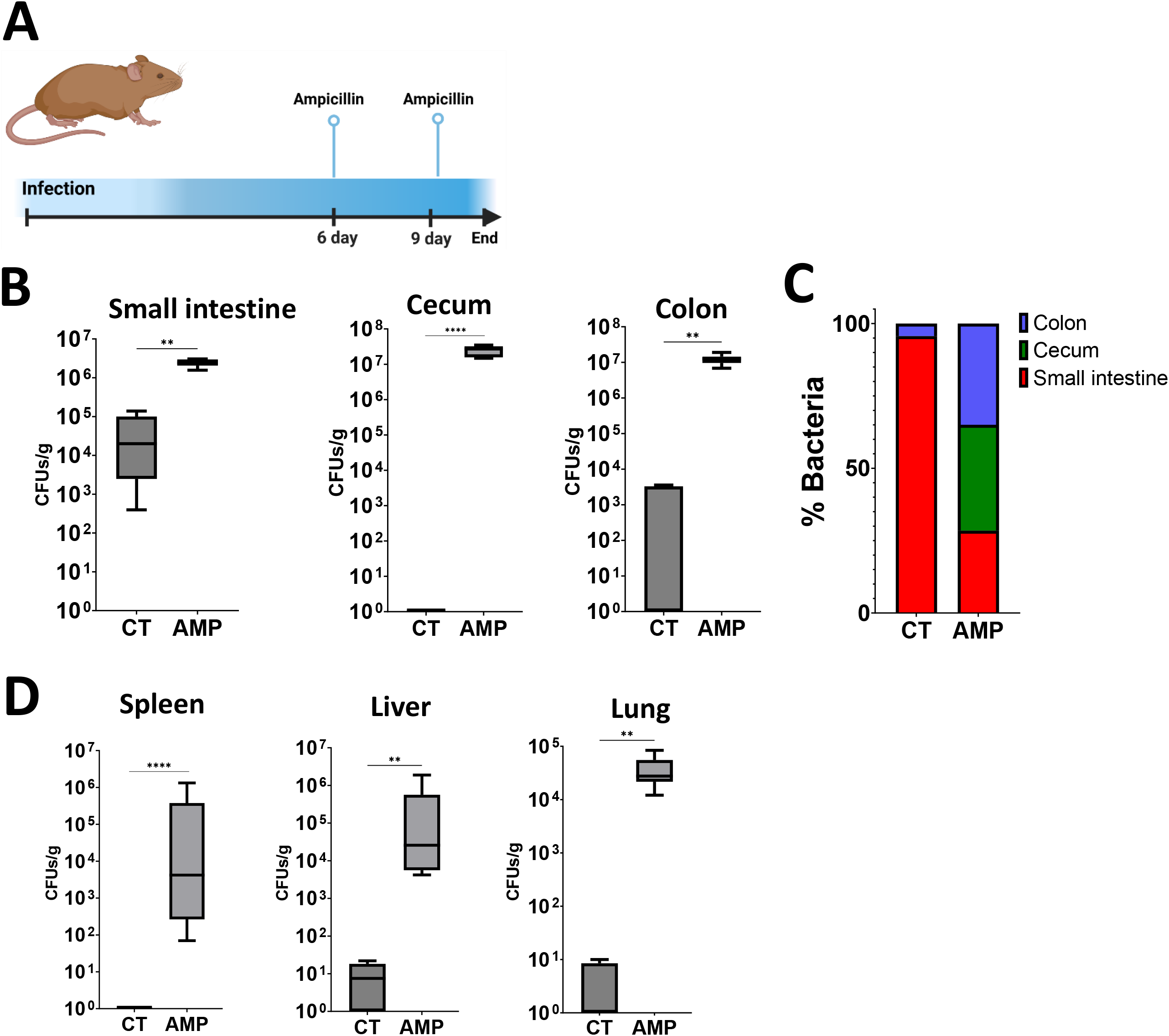
Antibiotic treatment triggers the dissemination of *K. pneumoniae* from the gastrointestinal tract. **A**. Six days after mice were colonised with strain Kp43816, mice were given two doses of ampicillin (i.p.) and 24 h after the last dose mice were euthanized and the bacterial loads were determined. **B**. CFUs per gr of small intestine, cecum, and colon of mice colonised with Kp43816 and treated with either vehicle solution (CT) or with ampicillin (AMP). **C**. Relative distribution of Kp43816 across the intestine sections of mice treated with vehicle solution (CT) or with ampicillin (AMP). **D**. Bacterial loads in the spleen, liver and lung of mice colonised with Kp43816 and treated with either vehicle solution (CT) or with ampicillin (AMP). In all panels, 6 mice were included in each group. Values are presented as the mean ± SD. ****P ≤ 0.0001; ** P ≤ 0.01 for the indicated comparisons determined using Mann-Whitney U test.

### The capsule and the type VI secretion system contribute to gastrointestinal tract colonisation

The fact that the CPS plays a crucial role in host-*K. pneumoniae* interface led us to examine the contribution of the CPS to gastrointestinal colonisation. There are reports showing that the CPS is necessary for gut colonisation of antibiotic pre-treated mice (22, 23). In our model, the *cps* mutant colonised the small intestine as well as the wild-type strain (Fig 6A). In contrast, the mutant colonized poorly the cecum and colon, suggesting that the CPS is necessary for the colonization of the large intestine. Further confirming this finding, distribution analysis of bacteria showed that the levels of the *cps* mutant were higher in the small intestine than in the large intestine (Fig 6B).

**Figure 6.**
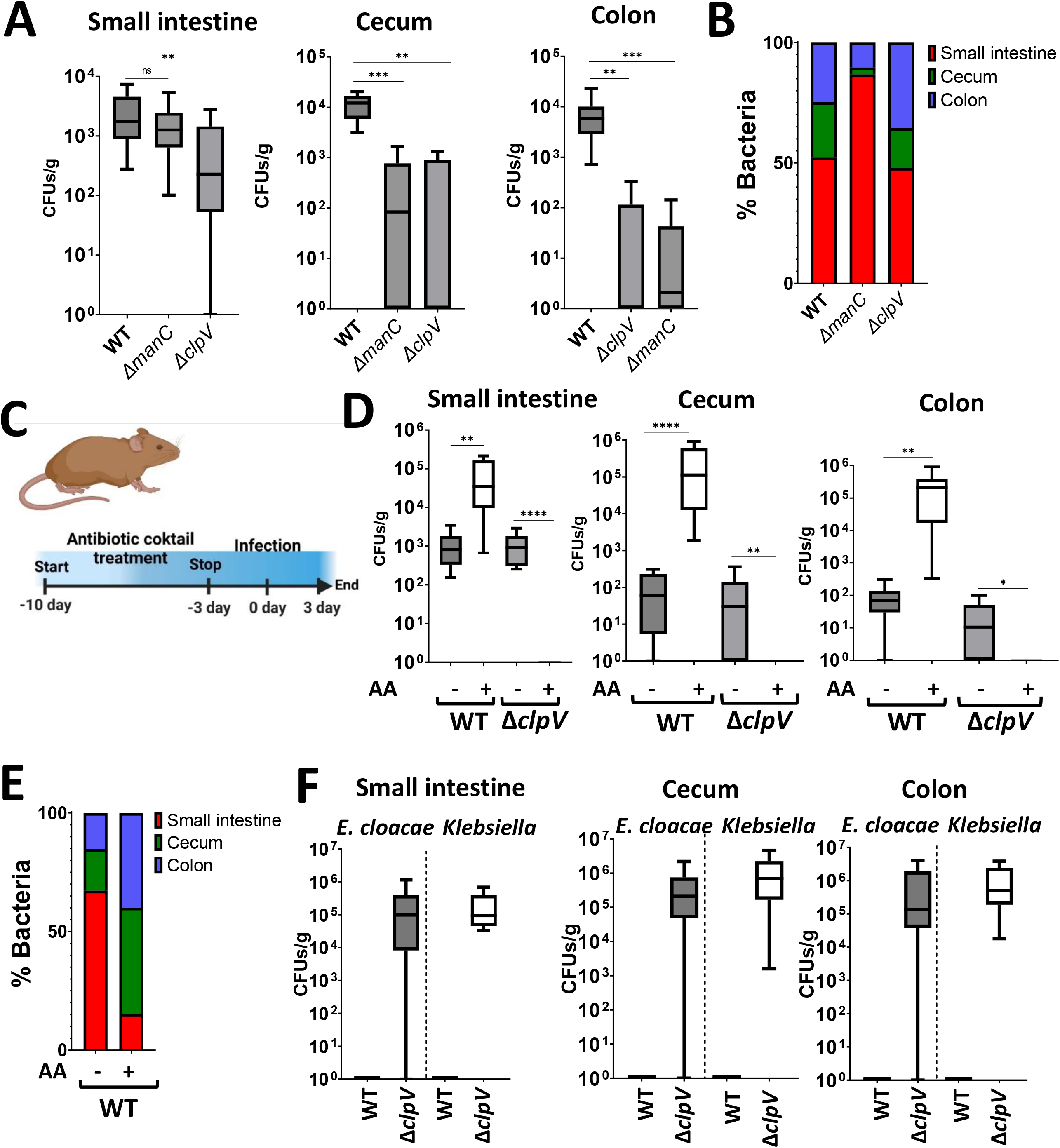
*K. pneumoniae* CPS and the T6SS are required for gut colonisation. **A**. CFUs per gr of small intestine, cecum, and colon of mice infected with Kp52145, the isogenic *cps* mutant (Δ*manC*) and the T6SS mutant (Δ*clpV*). 18-21 mice were included in each group in two independent experiments. **B**. Relative distribution of Kp52145, the isogenic cps mutant (Δ*manC*) and the T6SS mutant (Δ*clpV*) across the intestine sections. **C**. To determine the effect of the gut microbiome on the colonisation by the T6SS mutant, mice were pre-treated with an antibiotic cocktail which was stooped three days pre infection. Three days post infection, the bacterial burden in tissues was determined. **D**. CFUs per gr of small intestine, cecum, and colon of mice infected with Kp52145, and the T6SS mutant (Δ*clpV*) which were pre-treated or not with the antibiotic cocktail (AA). 10-16 mice were included in each group in two independent experiments. **E**. Relative distribution of Kp52145 across the intestine sections of mice pre-treated or not with the antibiotic cocktail (AA). **F**. CFUs per gr of small intestine, cecum, and colon of *E. cloacae* and *Klebsiella spp* only found in mice infected with the T6SS mutant (Δ*clpV*) and pre-treated with the antibiotic cocktail. In all panels, values are presented as the mean ± SD. ****P ≤ 0.0001; ***P ≤ 0.001** P ≤ 0.01; *P ≤ 0.05; ns P > 0.05 for the indicated comparisons determined using one way-ANOVA with Bonferroni contrast for multiple comparisons test.

We have recently demonstrated that *K. pneumoniae* exploits a T6SS for bacteria and fungi antagonism (24). Given the barrier imposed by the gut microbiome to *K. pneumoniae* gut colonisation, we sought to establish whether the T6SS is needed for the colonisation of the gastrointestinal tract. Mice were infected with the *clpV* mutant and the bacterial loads in the different sections of the gastrointestinal tract determined. ClpV is the AAA^+^ ATPase of the T6SS (25, 26), and in the *clpV* mutant background *K. pneumoniae* T6SS is not functional (24). The loads of the *clpV* mutant in the small intestine, cecum, and colon were significantly lower than those of the wild-type strain (Fig 4A), illustrating the need for a functional T6SS to colonise the gastrointestinal tract by *K. pneumoniae*. Interestingly, distribution analysis of the *clpV* mutant did not reveal any difference with the wild-type strain (Fig 6B), suggesting that the lack of function of the T6SS did not affect the distribution of the mutant across the gastrointestinal tract.

We next sought to ascertain whether a reduction in the gut microbiome before infection would mitigate the absence of a functional T6SS due to the limited competition exerted by the gut microbiome. Mice were pre-treated with an antibiotic cocktail for ten days before infection, and after three days without treatment, they were infected with the wild-type strain and the *clpV* mutant (Fig 6C). As we anticipated, there was a significant increase in the Kp52145 loads in the small intestine, cecum and colon in those mice pre-treated with antibiotics compared to the control ones (Fig 6D). Distribution analysis of Kp52145 distribution across the gastrointestinal tract showed a shift towards colonisation of the large intestine (Fig 6E). Only in antibiotic pre-treated mice, we observed dissemination of Kp52145 to the lungs and spleen (Fig S3). In contrast, the *clpV* mutant did not colonise the gastrointestinal tract of mice pre-treated with antibiotics (Fig 6D). This result may indicate that the T6SS is crucial for colonisation of the gastrointestinal tract even when the colonisation resistance imposed by the microbiota is disrupted. However, we noted that in those plates from the pre-treated mice infected with the *clpV* mutant appeared colonies that did not metabolize inositol, and other small colonies inositol positive. MALDI-TOF analysis revealed that the inositol negative bacteria were classified as *Enterobacter cloacae* (experiment 1 and 2), whereas the inositol positive bacteria were *K. variicola* (experiment 1*) and K. oxytoca* (experiment 2). These bacteria colonised in high levels only the small intestine, cecum, and colon of antibiotic pre-treated mice infected with the *clpV* mutant (Fig 6F). We reasoned that the antibiotic pre-treatment triggered a bloom of these bacteria present in low levels in the gut microbiome; however Kp52145 outcompeted them in a T6SS-dependent manner. To confirm this, we tested whether Kp52145 exerts antibacterial activity against these bacteria in a T6SS-dependent manner. Indeed, quantitative competition assays revealed that Kp52145 killed the *E. cloacae* and *Klebsiella spp* strains (Fig 7A). This was not the case when the prey strains were co-incubated with the *clpV* mutant (Fig 7A). To determine whether the *Klebsiella spp* strains outcompeted the *clpV* mutant in a T6SS-dependent manner, we constructed *K. oxytoca* and *K. variicola clpV* mutants. When the Kp52145 *clpV* mutant was co-incubated with any of the T6SS mutants there was no reduction in the recovery of the prey (Fig 7B).

**Figure 7.**
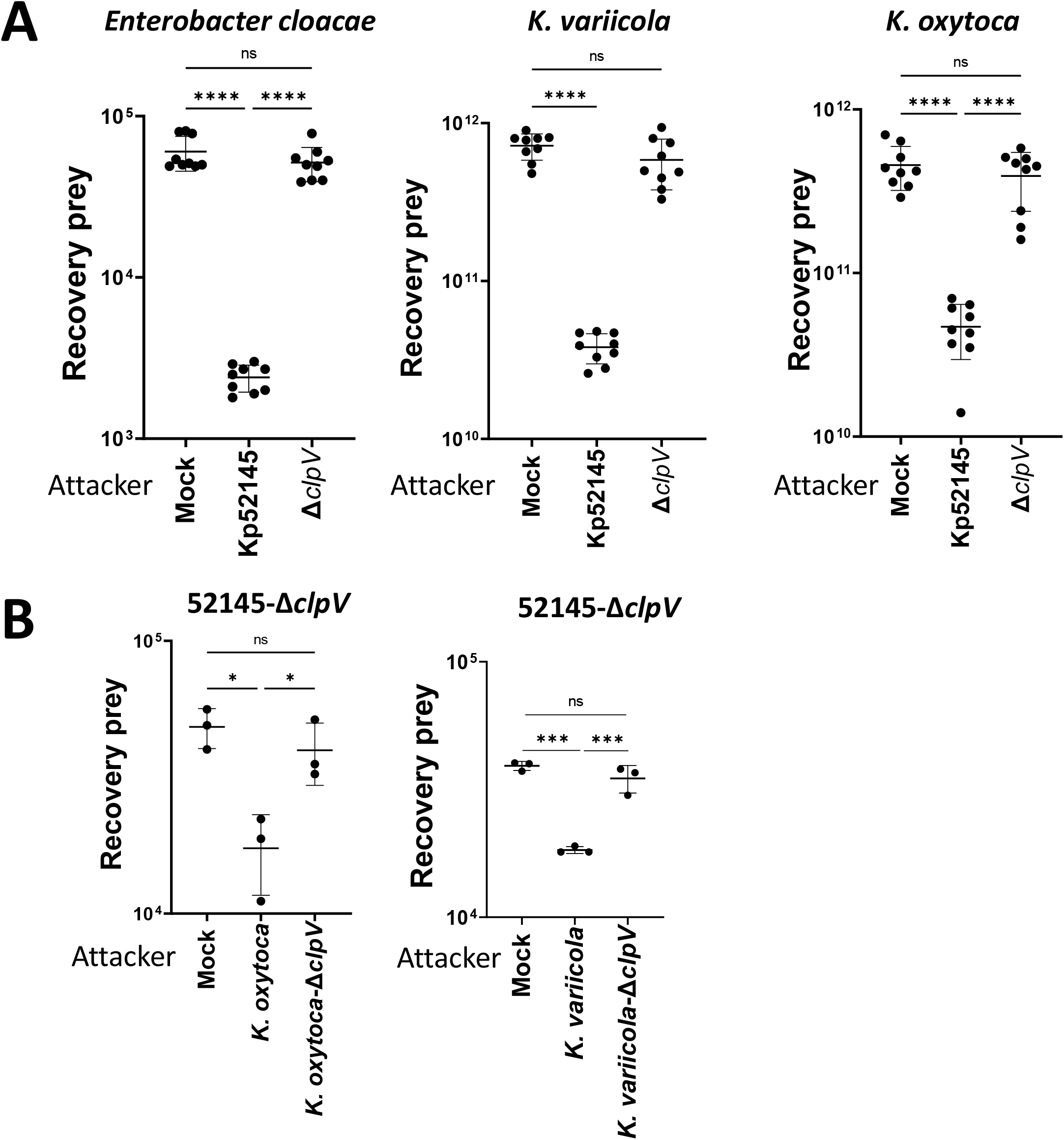
*K. pneumoniae* antagonises other *Enterobacteriaceae* of the gut microbiome in a T6SS-dependent manner. **A**. Bacterial killing mediated by Kp52145 and the T6SS mutant 52145-ΔclpV (Δ*clpV*) against *E. cloacae, K. oxytoca*, and *K. variicola*. Mock, PBS-treated prey. Number of recovered target cells following 6 h incubation in LB is indicated. **B**. Bacterial killing mediated by *K. oxytoca* and its isogeneic *clpV* mutant (*K. oxytoca*-Δ*clpV*), and by *K. variicola* and its *clpV* mutant (strain *K. variicola*-Δ*clpV*) against 52145-Δ*clpV* Mock, PBS-treated prey. Number of recovered target cells following 6 h incubation in LB is indicated. In all panels, the data are presented as means ± SD (n = 3). ****P ≤ 0.0001; ***P ≤ 0.001; *P ≤ 0.05; ns P > 0.05 for the indicated comparisons determined using one way-ANOVA with Bonferroni contrast for multiple comparisons test.

Altogether, our data demonstrate that the CPS is only necessary for the colonisation of the large intestine whereas the T6SS contributes to colonisation across the gastrointestinal tract. Our findings are consistent with the notion that *K. pneumoniae* exploits its T6SS to compete with the gut microbiome.

## DISCUSSION

In this work, we present a new model of gut colonisation by *K. pneumoniae* upon oral gavage that recapitulates key features of the asymptomatic human gastrointestinal tract colonisation by this pathogen. In our model, there is no need to disturb the microbiota of immunocompetent mice to achieve stable colonization without dissemination to other tissues. The fact that limited weight loss, histopathology, and inflammation were observed during colonisation indeed supports the notion that we are modelling asymptomatic colonisation without overt infection or disease. Importantly, our model also discriminates between *Klebsiella* strains with metastatic capacity and those without. We uncover that the small intestine is the primary site of colonisation by *K. pneumoniae*. Finally, we validate experimentally the clinical observation that antibiotic treatment favours *K. pneumoniae* dissemination from the gut to other tissues.

Previous studies used antibiotic treatment to disrupt the colonization barrier imposed by the gut microbiome to enable *K. pneumoniae* colonisation (11). As our work illustrates, this treatment shifts the distribution of *K. pneumoniae* within the gut increasing the colonisation of the large intestine and facilitating the dissemination to other tissues. Therefore, studies using antibiotic pre-treatment recapitulate an invasive gut infection rather than gut colonisation. Recently, an oral mouse model of infection also showed the ability of *K. pneumoniae* to colonize the gut without need of antibiotic pre-treatment (27). In contrast to our model, this model relies on the constant shedding of bacteria from the oropharynx, generating a continuous infection state of the mice exemplified by the high faecal shedding of bacteria (27). Although undoubtedly useful, this model does not recapitulate the asymptomatic gut colonization without upper airway infection characteristic of healthy human subjects.

Our findings establish that the primary site of colonization by *K. pneumoniae* is the small intestine without dissemination to other tissues. The microenvironment of the small intestine is remarkably different to that of the large intestine, being comparative more aerobic than other sections of the gastrointestinal tract, with lower density of microbiota than other sections, and with higher concentration of antimicrobial agents than other sections (28). These peptides create a harsh environment, limiting the colonization by enteric pathogens (29). However, work from our laboratory has uncovered a number of *Klebsiella* factors mediating antimicrobial peptides resistance (30-36), undoubtedly allowing *Klebsiella* to survive in this antimicrobial peptide rich microenvironment. The subsequent colonization of the colon after six days, although at lower levels than that the small intestine, suggests that the colonisation resistance to *K. pneumoniae* of the colon is higher than that of the small intestine. Although many factors may contribute to this barrier, our data indicate that the colon microbiome is a crucial one because its reduction increased dramatically *K. pneumoniae* bacterial loads. Notably, this effect was not that marked for the small intestine, suggesting that the microbiome of the small intestine confers limited resistance to *K. pneumoniae* colonization.

Unlike other *Enterobactericeae* such as *Salmonella typhimurium, Escherichia coli, Citrobacter rodentium* and *Vibrio cholerae* (37-43), *K. pneumoniae* colonisation was not associated with histopathology changes and with an acute inflammatory response in either the small or the large intestine. This finding does not contradict the clinical reports showing that *K. pneumoniae* is one of the pathogens associated with inflammatory bowel disease (44). This inflammatory condition triggers a strong dysbiosis, which in turn favours the bloom of pathogens, such as *Klebsiella*, already present within the gastrointestinal tract. Collectively, our results support the notion that *K. pneumoniae* colonisation of the gastrointestinal tract should be considered an example of stealth behaviour. Emerging evidence also indicates that *K. pneumoniae* stealthy strategies are crucial to overcome lung protective responses (45). A tantalizing hypothesis is that these stealth strategies were devoted initially to colonise the complex environment of the gastrointestinal tract in an asymptomatic manner but, when deployed to other tissues such as the lung, result in lethality. Future studies are warranted to validate this hypothesis by comparing the stealth strategies deployed by *Klebsiella* in the small intestine, the colon, and in the lung.

The studies done in mice pre-treated with antibiotics have revealed bacterial factors required for the infection of the gastrointestinal tract (11). These factors then play a role when the colonisation resistance is disrupted in immunocompetent mice. Therefore, there is a gap in our understanding of the factors employed by *K. pneumoniae* to counteract colonisation resistance. Here, we uncover that the CPS is necessary for the colonisation of the large intestine but dispensable for the colonisation of the small intestine, illustrating a hitherto unknown gastrointestinal microenvironment-dependent role of the CPS. The fact that the CPS is needed to colonise the large intestine of antibiotic pre-treated mice (22, 23) reveals the dual role of this virulence factor to overcome colonisation resistance and to colonise the gut when the barrier is disrupted. In the case of the former, there is evidence uncovering the role of the CPS as immunity factor against T6SS attack (46). Our findings add further weight to the notion that the CPS is the single most important factor governing *K. pneumoniae*-host interactions in vivo. Intriguingly, at present, it appears that the CPS is only dispensable in the case of urinary tract infections (47), questioning what makes the urinary tract so different compare to other tissues making the CPS dispensable.

We also tested the requirement of the T6SS in gastrointestinal colonisation. Previous work from our laboratory demonstrated the antimicrobial activity of *K. pneumoniae* T6SS against bacteria and fungi (24), making then plausible that the T6SS contributes to overcome the colonisation resistance imposed by the microbiome. Indeed, in our model the T6SS was required to colonise the small and large intestine. Furthermore, our data suggests that the primary role of the T6SS is to outcompete the gut microbiome. Similar results have been reported for other enteric pathogens such as *Shigella sonnei* and *S. typhimurium* (48, 49), illustrating the evolutionary conserved role of the T6SS to survive in the gut independently of the infection biology of the pathogen, a stealth coloniser like *K. pneumoniae* or invasive inflammatory pathogens such as *Shigella* and *Salmonella*.

Finally, it is worth discussing the translational opportunities offered by our model. Now it is possible to carry functional genomic studies to identify *K. pneumoniae* factors required to overcome colonisation resistance and to better understand the differences between metastatic *Klebsiella* and those strains that do not disseminate to other tissues. This work will provide a comprehensive understanding of the interface between *K. pneumoniae*, the gastrointestinal tract and the innate immune system, and the microbiome. Our model also allows the investigation of which factors, other infections or treatments for example, facilitate colonisation, or trigger the dissemination of *K. pneumoniae* from the gut. This knowledge is relevant to identify risks associated with *K. pneumoniae* invasive infections which are known to arise from gastrointestinal colonisation. Lastly, we envision our model will be an excellent platform to test therapeutics aiming to eliminate the asymptomatic colonisation of *K. pneumoniae*.

## METHODS

### Ethics statement

Experiments involving mice were approved by the Queen’s University Belfast’s Ethics Committee and conducted in accordance with the UK Home Office regulations (project licences PPL2778 and PPL2910) issued by the UK Home Office. Animals were randomized for interventions but researches processing the samples and analysing the data were aware which intervention group corresponded to which cohort of animals.

### Mice

All experiments were carried out in Queen’s University Biological Services Unit challenging 8-9 weeks old mice, using equal number of male and female animals. C57BL/6 mice were purchased from Charles River Laboratories and placed in sterile cages at their arrival during at least one week before experiment start for acclimation. Animals were supplied with food and water *ad libitum* and placed in individually ventilated cages when moved to the Biosafety Level 2 laboratory where the infection experiments were performed.

### Bacterial strains and growth conditions

Kp52145 is a *K. pneumoniae* clinical isolate (serotype O1:K2; sequence type ST66) previously described (13, 50). The capsule mutant strain, 52145-Δ*manC*, and the type VI secretion system mutant strain, 52145-Δ*clpV*, are isogenic strains of Kp52145 and they have been described previously (24, 35). NJST258-1 (serotype O1:K2; sequence type ST258); ATCC43816 (serotype O1:K2; sequence type ST493); and SGH10 (serotype K1; sequence type ST23) are also *K. pneumoniae* strains previously described (15-17). *K. oxytoca* MG2, *K. variicola* MG2, *E. cloacae* MG1 were isolated from the small intestine of mice infected with 52145-Δ*clpV* after antibiotic treatment.

All bacteria were grown in 5 mL Luria-Bertani (LB) medium at 37°C on an orbital shaker (180 rpm).

### Gut colonisation model

Overnight bacteria cultures were refreshed 1/10 into a new tube containing 4.5 mL of fresh LB media. After 2.5 hours of incubation on an orbital shaker (180 rpm) at 37°C, bacteria were pelleted (2500g, 20 minutes, RT) and suspended in PBS. The suspension was adjusted to an OD_600_ of 1.0 to obtain a concentration of 5 × 10^8^ CFU/ml.

At the start of the gut colonisation experiments, animals were administered 200 μl of a 0.2M solution of sodium bicarbonate by oral gavage. 5 min later, mice were infected with 100 μl of the desired inoculum. Mice were weighted and checked daily to detect any sign of distress. Animals showing moderate signs of distress (inactivity, separation from the group, hunched posture and/or body weight loss >10% in two consecutive days or >20% in 24 hours) were euthanized by approved schedule 1 protocol. Lungs and spleens were plated to asses dissemination of the bacteria.

### Quantification of *K. pneumoniae* burden

Small intestine, cecum, colon, faecal samples, lungs, liver and spleen from mice were weighted, immersed in 1 ml of PBS and processed for bacterial quantification. In the case of gut samples, intestinal content was previously removed by gently pushing it from one end of the gut and out the other end. Samples were homogenized by mechanical disruption with a sterile plastic loop (faecal samples), or with a Precellys Evolution tissue homogenizer (Bertin Instruments) using 1.4 mm ceramic (zirconium oxide) beads at 4,500 rpm for 7 cycles of 10 seconds, with a 10 second-pause between each cycle (small intestine, cecum, colon, lungs, liver and spleen).

Homogenates were serially diluted in sterile PBS and plated onto *Salmonella-Shigella* agar (Sigma-Aldrich) (lungs, liver and spleen), or SCAI medium [Simmons Citrate (Sigma-Aldrich) agar with 1% inositol] (small intestine, cecum, colon and faecal samples), for selective identification of *K. pneumoniae* colonies. Colonies were enumerated after overnight incubation at 37°C in case of *Salmonella-Shigella* agar, or after 72 hours incubation in case of the SCAI plates.

### Histology analysis

Small intestine and colon were excised from the animals and their contents removed as previously described. Further cleaning was performed by gently passing through the lumen 1 ml of 4% paraformaldehyde (PFA). Organs were fixed 24 hours in an excess volume of 4% PFA and placed in PBS until being processed.

For processing, samples dehydrated, embedded in paraffin and 10 μm sections were prepared. Haematoxylin and eosin (H&E) staining of the sections was performed according to standard protocols. Images of the H&E stained slides were obtained with a Eclipse 80i microscope (Nikon) using a 20x/0.8NA objective, with digital images acquired with the NIS-Elements software (Nikon).

Blinded evaluation was performed by a researcher of the laboratory using the histopathology scoring indicated in Table S1.

### RNA isolation and RT-qPCR

Content-cleaned small intestine, cecum and colon were collected in RNA stabilizing solution and homogenized with a handheld homogenizer in 1 ml of TRIzol reagent (Ambion), for total RNA extraction according to the manufacturer’s instructions. Extracted RNA was treated with DNase I (Roche), precipitated with sodium acetate and ethanol, and quantified using a Nanovue Plus spectrophotometer (GE Healthcare Life Sciences). cDNA was generated by retrotranscription of 1 g of total RNA using M-MLV reverse transcriptase (Invitrogen) and random primers (Invitrogen).

Ten nanograms of cDNA were used as a template in a 5 μl reaction mixture from a KAPA SYBR FAST qPCR kit (Kapa Biosystems). Primers used are listed in Table S2. RT-qPCR was performed using a Rotor-Gene Q (Qiagen) with the following thermocycling conditions: 95°C for 3 min for hot-start polymerase activation, followed by 40 cycles of 95°C for 5 s and 60°C for 20 s. Fluorescence of SYBR green dye was measured at 510 nm. Relative quantities of mRNAs were obtained using the ΔΔC_T_ method by using hypoxanthine phosphoribosyltransferase 1 (*hprt*) gene normalization.

### Microbiome analysis

Small intestine and colon faecal contents were collected in sterile tubes and stored at −80°C until processed. Microbial DNA was extracted using a QIAamp PowerFecal Pro DNA Kit (Qiagen), according to manufacturer’s instructions, using a Precellys Evolution tissue homogenizer (Bertin Instruments) at 5,000 rpm for 2 cycles of 30 seconds, with a 30 seconds-pause between them, to homogenize the samples. Extracted DNA was quantified using a Nanovue Plus spectrophotometer (GE Healthcare Life Sciences).

Processing and sequencing of the DNA samples was performed by the Genomics Core Technology Unit of Queen’s University Belfast. Libraries were constructed in batches, quantified and run on the Illumina Miseq V2 Platform with a read length of 500 bp (2 × 250 bp) and a read depth of 100.000 reads/sample. 16S amplicon PCR of the V3-V4 region was performed using 341F (CCTACGGGNGGCWGCAG) and 806R (GGACTACHVGGGTWTCTAAT) primers.

Amplicon identification, quantification and analyses were carried out in R 4.1.1 using different packages. Firstly, a quality assessment of the sequences was performed using FASTQC software (https://www.bioinformatics.babraham.ac.uk/projects/fastqc/), and then a DADA2 (51) version 1.24 pipeline was used to generate the amplicon sequence variants (ASVs). In brief it consisted in a first stage were sequences were inspected visually prior to apply the filterAndTrim function (trimming 13 bases from each sequence, removing sequences containing Ns and applying a filter for the Expected Errors (maxEE) of 2 and 6 for each of the reads) retaining on average 76% of the reads. Then error models were learnt and applied to denoise and merge reads, using default parameters prior to the removal of chimeric sequences. SILVA v138 database was used for the taxonomic assignation using a minimum bootstrap confidence of 80, identifying a total of 2988 ASVs that were transformed into a phyloseq object for further analyses and further filtered down by removing sequences assigned to mitochondria, and those without phylum or class. Further filtering based on prevalence and abundance resulted in a total of 819 ASVs. Alpha and beta diversity were analysed using Phyloseq v1.40 (52), and a differential abundance analysis was carried out using Deseq2 v1.36.0 using a statistical significance threshold of adj. p-value = 0.05. Data are available at NCBI SRA (bioproject PRJNA880484).

### Post-infection antibiotic treatment

C57BL/6 mice were infected with 1×10^7^ CFUs of the ampicillin-resistant *K. pneumoniae* strain ATCC43186, as described before. On day 6 and 9 post inoculation, they received a 500 mg/kg of ampicillin sodium salt (Sigma-Aldrich), or vehicle (sterile water) intraperitoneally. 24 hours after the second dose, animals were euthanized and organs collected for bacterial burden quantification as previously described.

### Microbiota depletion

C57BL/6 mice were treated for 10 days with a mix of broad-spectrum antibiotics in the drinking water (ampicillin 1g/l, neomycin sulfate 1 g/l, metronidazole 1 g/l, and vancomycin 0.5 g/l) (Sigma-Aldrich) (Brown et al., 2017). Antibiotic therapy was stopped 3 days prior the infection, replacing it with normal, antibiotic free, drinking water. Mice were infected with 1×10^7^ CFUs of either Kp52145 or 52145-Δ*clpV* strains via oral gavage as described before. Animals were euthanized 72 hours post-inoculation, and organs processed for bacterial quantification as previously described.

### Identification of bacterial colonies by MALDI

When required, identification of bacterial colonies was performed using a VITEK MS instrument (BioMerieux, France), in accordance with manufacturer’s instructions. Briefly, fresh cultures were prepared on appropriate media from which a single bacterial colony, or part of a colony, was transferred to the target slide as a smear. Then, 1 μl of matrix (Vitek MS-CHCA) was added and air dried. Following insertion of the loaded slide into the VITEK MS, operating with software version 1.6.0, spectra were generated by the instrument from the bacterial suspensions and compared to reference spectra in the database to provide an identification. Each identification was reported with a score, expressed as a percentage, indicating the degree of confidence in that result. Scores of >90% were considered decisive.

### Antibacterial competition assay

Attacker *K. pneumoniae* strains and preys were grown until mid-exponential phase, collected by centrifugation and resuspended in PBS to an OD_600_ of 1.2. Attacker and preys were mixed at 10:1 ratio by volume, and 100 μl of the mixture was spotted in a pre-warmed agar plate at 37°C. The contact spot was incubated for 6 hours. Recovered cells were plated out on antibiotic selective media and viable cells were reported as recovery target cell representing the CFU per ml of the recovered prey after co-culture. *K. variicola, K. oxytoca* and *E. cloacae* were selected by plating on 25 μg/ml carbenicillin, and 52145-Δ*clpV* was selected by plating on LB agar containing 3 μg/ml sodium tellurite. All experiments were carried out with triplicate samples on at least three independent occasions.

### Construction of *clpV* mutants

Strains were grown in Luria-Bertani media, supplemented with appropriate antibiotics at the following concentrations: gentamycin (30 μg/mL), chloramphenicol (30 μg/mL), kanamycin (100 μg/mL) and diaminopimelic acid (DAP, 300 μM). Bacterial genomic DNA was extracted with PureLink Genomic DNA Kit (Life Technologies) as per manufacturer’s instructions. DNA fragments targeting the *clpV* gene were amplified from genomic DNA using KAPA HiFi DNA polymerase (Roche). Primers were designed from conserved regions identified by comparison of the *clpV* gene from the available *K. variicola* or *K. oxytoca* genomes on NCBI Genbank (accessed 01/07/22 and 04/02/2022, respectively). Primers are shown in Table S2.

To construct a *K. oxytoca clpV* mutant, an internal fragment of *clpV* was amplified by PCR using primers pKNOCK-Koxy-clpV-F3 and pKNOCK-Koxy-clpV-R3. The PCR product was gel purified and it was cloned into the pKNOCK-cm suicide vector (53), cut with BamHI (New England Biolabs), by homologous alignment cloning (54) to generate pKNOCKClpV_oxy_. This plasmid was transformed into the *E. coli* DAP auxotroph β2163 (55) that then mobilized the plasmid to *K. oxytoca* MG2. A mutant in which pKNOCKClpV_oxy_ was inserted by homologous recombination into the *clpV* locus was confirmed by PCR using primers pKNOCK-Koxy-clpV-sc-F and pKNOCK-Koxy-clpV-sc-R, and named *K. oxytoca-ΔclpV*.

To construct a *K. variicola clpV* mutant, we used the λ-Red-mediated homologous recombination as described previously (56). Homologous regions up- and down-stream of *clpV* were amplified by PCR using primers 6144-clpV.2 and 6144-clpV.3, and 6144-clpV.4 and 6144-clpV.5 respectively, and sewn together with the kanamycin cassette amplified from pKD4 (57) using primers cm.3a and cm.4a. KAPA HiFi was used as polymerase. PCR products were amplified with using primers 6144-clpV.2 and 6144-clpV.5, and the PCR fragment was purified with QIAGEN MinElute kit. Strain *K. variicola* MG2 was transformed with pKOBEG-gent encoding λ phage redγβα operon (58) expressed under the control of the arabinose-inducible pBAD promoter. Briefly, *K. variicola* were grown overnight in LB and then sub-cultured 1:100 in 200 mL of LB supplemented with gentamycin and 0.2% arabinose. Cells were grown at 37 °C until reaching an OD_600_ of 0.4. Cells were cooled on ice for 30 minutes then pelleted and resuspended in ice-cold 10% glycerol and incubated on ice for 1 hour. Cells were washed three times with ice-cold 10% glycerol and finally suspended in 200 μl of 10% glycerol. The cells were electroporated with 1 ug 3-way-PCR DNA in a 2mm cuvette with 2.5 kV, 200 Ω and 25 μF. Cells were recovered in 1 mL of SOC media (2% tryptone, 0.5% yeast extract, 10 mM NaCl, 2.5 mM KCl, 10 mM MgCl_2_, 10 mM MgSO_4_, and 20 mM glucose) at 37 C for 2 hours. Transformed cells were selected by plating on LB agar supplemented with kanamycin. The replacement of the wild-type allele by the mutant was confirmed by PCR using primers 6144-clpV.1 and 6144-clpV.6, and the mutant named *K. variicola-ΔclpV*. pKOBEG-gent plasmid was cured from the mutant strain by overnight growth at 37°C.

### Quantification and statistical analysis

Statistical analyses were performed using one-way analysis of variance (ANOVA) with Bonferroni corrections, the one-tailed t test, or, when the requirements were not met, the Mann-Whitney U test. p values of <0.05 were considered statistically significant. Normality and equal variance assumptions were tested with the Kolmogorov-Smirnov test and the Brown-Forsythe test, respectively. All analyses were performed using GraphPad Prism for Windows (version 9.4.1) software.

## ACKNOWLEDGEMENTS

We thank the members of the J.A.B. laboratory for their thoughtful discussions and support with this project. This work was supported by the Trond Mohn Foundation (contract TMF2019TMT03), and by Biotechnology and Biological Sciences Research Council (BBSRC, BB/V007939/1, BBW510682/1), and Medical Research Council (MRC, MR/V032496/1) funds to J.A.B.

## REFERENCES

1. Antimicrobial Resistance C. 2022. Global burden of bacterial antimicrobial resistance in 2019: a systematic analysis. Lancet 399:629–655.

2. Russo TA, Marr CM. 2019. Hypervirulent Klebsiella pneumoniae. Clinical microbiology reviews 32:10.1128/CMR.00001-19. Print 2019 Jun 19.

3. Gorrie CL, Mirceta M, Wick RR, Edwards DJ, Thomson NR, Strugnell RA, Pratt NF, Garlick JS, Watson KM, Pilcher DV, McGloughlin SA, Spelman DW, Jenney AWJ, Holt KE. 2017. Gastrointestinal Carriage Is a Major Reservoir of Klebsiella pneumoniae Infection in Intensive Care Patients. Clinical infectious diseases: an official publication of the Infectious Diseases Society of America 65:208–215.

4. Martin RM, Cao J, Brisse S, Passet V, Wu W, Zhao L, Malani PN, Rao K, Bachman MA. 2016. Molecular Epidemiology of Colonizing and Infecting Isolates of Klebsiella pneumoniae. mSphere 1:10.1128/mSphere.00261-16. eCollection 2016 Sep-Oct.

5. Gu D, Dong N, Zheng Z, Lin D, Huang M, Wang L, Chan EW, Shu L, Yu J, Zhang R, Chen S. 2018. A fatal outbreak of ST11 carbapenem-resistant hypervirulent Klebsiella pneumoniae in a Chinese hospital: a molecular epidemiological study. The LancetInfectious diseases 18:37–46.

6. Yao H, Qin S, Chen S, Shen J, Du XD. 2018. Emergence of carbapenem-resistant hypervirulent Klebsiella pneumoniae. The LancetInfectious diseases 18:25-3099(17)30628-X. Epub 2017 Nov 1.

7. Zhang Y, Zeng J, Liu W, Zhao F, Hu Z, Zhao C, Wang Q, Wang X, Chen H, Li H, Zhang F, Li S, Cao B, Wang H. 2015. Emergence of a hypervirulent carbapenem-resistant Klebsiella pneumoniae isolate from clinical infections in China. The Journal of infection 71:553–560.

8. Lam MMC, Wyres KL, Wick RR, Judd LM, Fostervold A, Holt KE, Lohr IH. 2019. Convergence of virulence and MDR in a single plasmid vector in MDR Klebsiella pneumoniae ST15. J Antimicrob Chemother 74:1218–1222.

9. Raffelsberger N, Hetland MAK, Svendsen K, Smabrekke L, Lohr IH, Andreassen LLE, Brisse S, Holt KE, Sundsfjord A, Samuelsen O, Gravningen K. 2021. Gastrointestinal carriage of Klebsiella pneumoniae in a general adult population: a cross-sectional study of risk factors and bacterial genomic diversity. Gut Microbes 13:1939599.

10. Donskey CJ. 2004. The role of the intestinal tract as a reservoir and source for transmission of nosocomial pathogens. Clin Infect Dis 39:219–26.

11. Joseph L, Merciecca T, Forestier C, Balestrino D, Miquel S. 2021. From Klebsiella pneumoniae Colonization to Dissemination: An Overview of Studies Implementing Murine Models. Microorganisms 9.

12. Holt KE, Wertheim H, Zadoks RN, Baker S, Whitehouse CA, Dance D, Jenney A, Connor TR, Hsu LY, Severin J, Brisse S, Cao H, Wilksch J, Gorrie C, Schultz MB, Edwards DJ, Nguyen KV, Nguyen TV, Dao TT, Mensink M, Minh VL, Nhu NT, Schultsz C, Kuntaman K, Newton PN, Moore CE, Strugnell RA, Thomson NR. 2015. Genomic analysis of diversity, population structure, virulence, and antimicrobial resistance in Klebsiella pneumoniae, an urgent threat to public health. Proceedings of the National Academy of Sciences of the United States of America 112:E3574–81.

13. Lery LM, Frangeul L, Tomas A, Passet V, Almeida AS, Bialek-Davenet S, Barbe V, Bengoechea JA, Sansonetti P, Brisse S, Tournebize R. 2014. Comparative analysis of Klebsiella pneumoniae genomes identifies a phospholipase D family protein as a novel virulence factor. BMC biology 12:41-7007-12-41.

14. Van Kregten E, Westerdaal NA, Willers JM. 1984. New, simple medium for selective recovery of Klebsiella pneumoniae and Klebsiella oxytoca from human feces. J Clin Microbiol 20:936–41.

15. Budnick JA, Bina XR, Bina JE. 2021. Complete Genome Sequence of Klebsiella pneumoniae Strain ATCC 43816. Microbiol Resour Announc 10.

16. Deleo FR, Chen L, Porcella SF, Martens CA, Kobayashi SD, Porter AR, Chavda KD, Jacobs MR, Mathema B, Olsen RJ, Bonomo RA, Musser JM, Kreiswirth BN. 2014. Molecular dissection of the evolution of carbapenem-resistant multilocus sequence type 258 Klebsiella pneumoniae. Proceedings of the National Academy of Sciences of the United States of America 111:4988–4993.

17. Lam MMC, Wick RR, Wyres KL, Gorrie CL, Judd LM, Jenney AWJ, Brisse S, Holt KE. 2018. Genetic diversity, mobilisation and spread of the yersiniabactin-encoding mobile element ICEKp in Klebsiella pneumoniae populations. Microbial genomics doi:10.1099/mgen.0.000196 [doi].

18. Winter SE, Winter MG, Xavier MN, Thiennimitr P, Poon V, Keestra AM, Laughlin RC, Gomez G, Wu J, Lawhon SD, Popova IE, Parikh SJ, Adams LG, Tsolis RM, Stewart VJ, Baumler AJ. 2013. Host-derived nitrate boosts growth of E. coli in the inflamed gut. Science 339:708–11.

19. Sands BE. 2015. Biomarkers of inflammation in inflammatory bowel disease. Gastroenterology 149:1275-1285. e2.

20. Edwards JA, Tan N, Toussaint N, Ou P, Mueller C, Stanek A, Zinsou V, Roudnitsky S, Sagal M, Dresner L, Schwartzman A, Huan C. 2020. Role of regenerating islet-derived proteins in inflammatory bowel disease. World J Gastroenterol 26:2702–2714.

21. Shin JH, Seeley RJ. 2019. Reg3 Proteins as Gut Hormones? Endocrinology 160:1506–1514.

22. Favre-Bonte S, Licht TR, Forestier C, Krogfelt KA. 1999. Klebsiella pneumoniae capsule expression is necessary for colonization of large intestines of streptomycin-treated mice. Infect Immun 67:6152–6.

23. Tan YH, Chen Y, Chu WHW, Sham LT, Gan YH. 2020. Cell envelope defects of different capsule-null mutants in K1 hypervirulent Klebsiella pneumoniae can affect bacterial pathogenesis. Mol Microbiol 113:889–905.

24. Storey D, McNally A, Astrand M, Sa-Pessoa Graca Santos J, Rodriguez-Escudero I, Elmore B, Palacios L, Marshall H, Hobley L, Molina M, Cid VJ, Salminen TA, Bengoechea JA. 2020. Klebsiella pneumoniae type VI secretion system-mediated microbial competition is PhoPQ controlled and reactive oxygen species dependent. PLoS Pathog 16:e1007969.

25. Cherrak Y, Flaugnatti N, Durand E, Journet L, Cascales E. 2019. Structure and Activity of the Type VI Secretion System. Microbiol Spectr 7.

26. Bonemann G, Pietrosiuk A, Diemand A, Zentgraf H, Mogk A. 2009. Remodelling of VipA/VipB tubules by ClpV-mediated threading is crucial for type VI protein secretion. EMBO J 28:315–25.

27. Young TM, Bray AS, Nagpal RK, Caudell DL, Yadav H, Zafar MA. 2020. Animal Model To Study Klebsiella pneumoniae Gastrointestinal Colonization and Host-to-Host Transmission. Infect Immun 88.

28. Donaldson GP, Lee SM, Mazmanian SK. 2016. Gut biogeography of the bacterial microbiota. Nat Rev Microbiol 14:20–32.

29. Ostaff MJ, Stange EF, Wehkamp J. 2013. Antimicrobial peptides and gut microbiota in homeostasis and pathology. EMBO Mol Med 5:1465–83.

30. Llobet E, March C, Gimenez P, Bengoechea JA. 2009. Klebsiella pneumoniae OmpA confers resistance to antimicrobial peptides. Antimicrobial Agents and Chemotherapy 53:298–302.

31. Llobet E, Martinez-Moliner V, Moranta D, Dahlstrom KM, Regueiro V, Tomas A, Cano V, Perez-Gutierrez C, Frank CG, Fernandez-Carrasco H, Insua JL, Salminen TA, Garmendia J, Bengoechea JA. 2015. Deciphering tissue-induced Klebsiella pneumoniae lipid A structure. Proceedings of the National Academy of Sciences of the United States of America 112:E6369–78.

32. Llobet E, Tomas JM, Bengoechea JA. 2008. Capsule polysaccharide is a bacterial decoy for antimicrobial peptides. Microbiology (Reading, England) 154:3877–3886.

33. Llobet E, Campos MA, Gimenez P, Moranta D, Bengoechea JA. 2011. Analysis of the networks controlling the antimicrobial-peptide-dependent induction of Klebsiella pneumoniae virulence factors. Infection and immunity 79:3718–3732.

34. Padilla E, Llobet E, Domenech-Sanchez A, Martinez-Martinez L, Bengoechea JA, Alberti S. 2010. Klebsiella pneumoniae AcrAB efflux pump contributes to antimicrobial resistance and virulence. Antimicrobial Agents and Chemotherapy 54:177–183.

35. Kidd TJ, Mills G, Sa-Pessoa J, Dumigan A, Frank CG, Insua JL, Ingram R, Hobley L, Bengoechea JA. 2017. A Klebsiella pneumoniae antibiotic resistance mechanism that subdues host defences and promotes virulence. EMBO molecular medicine 9:430–447.

36. Campos MA, Vargas MA, Regueiro V, Llompart CM, Alberti S, Bengoechea JA. 2004. Capsule polysaccharide mediates bacterial resistance to antimicrobial peptides. Infection and immunity 72:7107–7114.

37. Mullineaux-Sanders C, Sanchez-Garrido J, Hopkins EGD, Shenoy AR, Barry R, Frankel G. 2019. Citrobacter rodentium-host-microbiota interactions: immunity, bioenergetics and metabolism. Nat Rev Microbiol 17:701–715.

38. Qadri F, Bhuiyan TR, Dutta KK, Raqib R, Alam MS, Alam NH, Svennerholm A-M, Mathan MM. 2004. Acute dehydrating disease caused by <em>Vibrio cholerae</em> serogroups O1 and O139 induce increases in innate cells and inflammatory mediators at the mucosal surface of the gut. Gut 53:62–69.

39. Millet YA, Alvarez D, Ringgaard S, von Andrian UH, Davis BM, Waldor MK. 2014. Insights into Vibrio cholerae intestinal colonization from monitoring fluorescently labeled bacteria. PLoS Pathog 10:e1004405.

40. Kaper JB, Nataro JP, Mobley HL. 2004. Pathogenic Escherichia coli. Nat Rev Microbiol 2:123–40.

41. Santos RL, Raffatellu M, Bevins CL, Adams LG, Tukel C, Tsolis RM, Baumler AJ. 2009. Life in the inflamed intestine, Salmonella style. Trends Microbiol 17:498–506.

42. Barthel M, Hapfelmeier S, Quintanilla-Martinez L, Kremer M, Rohde M, Hogardt M, Pfeffer K, Russmann H, Hardt WD. 2003. Pretreatment of mice with streptomycin provides a Salmonella enterica serovar Typhimurium colitis model that allows analysis of both pathogen and host. Infect Immun 71:2839–58.

43. Coburn B, Li Y, Owen D, Vallance BA, Finlay BB. 2005. Salmonella enterica serovar Typhimurium pathogenicity island 2 is necessary for complete virulence in a mouse model of infectious enterocolitis. Infect Immun 73:3219–27.

44. Federici S, Kredo-Russo S, Valdes-Mas R, Kviatcovsky D, Weinstock E, Matiuhin Y, Silberberg Y, Atarashi K, Furuichi M, Oka A, Liu B, Fibelman M, Weiner IN, Khabra E, Cullin N, Ben-Yishai N, Inbar D, Ben-David H, Nicenboim J, Kowalsman N, Lieb W, Kario E, Cohen T, Geffen YF, Zelcbuch L, Cohen A, Rappo U, Gahali-Sass I, Golembo M, Lev V, Dori-Bachash M, Shapiro H, Moresi C, Cuevas-Sierra A, Mohapatra G, Kern L, Zheng D, Nobs SP, Suez J, Stettner N, Harmelin A, Zak N, Puttagunta S, Bassan M, Honda K, Sokol H, Bang C, Franke A, Schramm C, Maharshak N, et al. 2022. Targeted suppression of human IBD-associated gut microbiota commensals by phage consortia for treatment of intestinal inflammation. Cell 185:2879–2898 e24.

45. Bengoechea JA, Sa Pessoa J. 2019. Klebsiella pneumoniae infection biology: living to counteract host defences. FEMS microbiology reviews 43:123–144.

46. Flaugnatti N, Isaac S, Lemos Rocha LF, Stutzmann S, Rendueles O, Stoudmann C, Vesel N, Garcia-Garcera M, Buffet A, Sana TG, Rocha EPC, Blokesch M. 2021. Human commensal gut Proteobacteria withstand type VI secretion attacks through immunity protein-independent mechanisms. Nat Commun 12:5751.

47. Ernst CM, Braxton JR, Rodriguez-Osorio CA, Zagieboylo AP, Li L, Pironti A, Manson AL, Nair AV, Benson M, Cummins K, Clatworthy AE, Earl AM, Cosimi LA, Hung DT. 2020. Adaptive evolution of virulence and persistence in carbapenem-resistant Klebsiella pneumoniae. Nat Med 26:705–711.

48. Anderson MC, Vonaesch P, Saffarian A, Marteyn BS, Sansonetti PJ. 2017. Shigella sonnei Encodes a Functional T6SS Used for Interbacterial Competition and Niche Occupancy. Cell Host Microbe 21:769–776 e3.

49. Sana TG, Flaugnatti N, Lugo KA, Lam LH, Jacobson A, Baylot V, Durand E, Journet L, Cascales E, Monack DM. 2016. Salmonella Typhimurium utilizes a T6SS-mediated antibacterial weapon to establish in the host gut. Proc Natl Acad Sci U S A 113:E5044–51.

50. Nassif X, Fournier JM, Arondel J, Sansonetti PJ. 1989. Mucoid phenotype of Klebsiella pneumoniae is a plasmid-encoded virulence factor. Infection and immunity 57:546–552.

51. Callahan BJ, McMurdie PJ, Rosen MJ, Han AW, Johnson AJ, Holmes SP. 2016. DADA2: High-resolution sample inference from Illumina amplicon data. Nat Methods 13:581–3.

52. McMurdie PJ, Holmes S. 2013. phyloseq: an R package for reproducible interactive analysis and graphics of microbiome census data. PLoS One 8:e61217.

53. Alexeyev MF. 1999. The pKNOCK series of broad-host-range mobilizable suicide vectors for gene knockout and targeted DNA insertion into the chromosome of gram-negative bacteria. BioTechniques 26:824-6, 828.

54. Tan L, Strong EJ, Woods K, West NP. 2018. Homologous alignment cloning: a rapid, flexible and highly efficient general molecular cloning method. PeerJ 6:e5146.

55. Demarre G, Guerout AM, Matsumoto-Mashimo C, Rowe-Magnus DA, Marliere P, Mazel D. 2005. A new family of mobilizable suicide plasmids based on broad host range R388 plasmid (IncW) and RP4 plasmid (IncPalpha) conjugative machineries and their cognate Escherichia coli host strains. Research in microbiology 156:245–255.

56. Hancock SJ, Phan MD, Luo Z, Lo AW, Peters KM, Nhu NTK, Forde BM, Whitfield J, Yang J, Strugnell RA, Paterson DL, Walsh TR, Kobe B, Beatson SA, Schembri MA. 2020. Comprehensive analysis of IncC plasmid conjugation identifies a crucial role for the transcriptional regulator AcaB. Nat Microbiol 5:1340–1348.

57. Datsenko KA, Wanner BL. 2000. One-step inactivation of chromosomal genes in Escherichia coli K-12 using PCR products. Proceedings of the National Academy of Sciences of the United States of America 97:6640–6645.

58. Balestrino D, Haagensen JA, Rich C, Forestier C. 2005. Characterization of type 2 quorum sensing in Klebsiella pneumoniae and relationship with biofilm formation. J Bacteriol 187:2870–80.

